# Subcortical magnocellular visual system facilities object recognition by processing topological property

**DOI:** 10.1101/2020.01.04.894725

**Authors:** Wenbo Wang, Tiangang Zhou, Yan Zhuo, Lin Chen, Yan Huang

## Abstract

The Magnocellular (M) visual pathway is known as a fast route to convey coarse information and facilitates object recognition by initiating top-down processes. It is unclear what exact properties M pathway conveys to accelerate visual object processing. Previous studies suggest that visual systems are highly sensitive to the perception of topological property (TP), which remains unchanged under various shape changes, and the TP is probably processed through a fast subcortical pathway. Here we hypothesize that a subcortical M system contributes to the fast object recognition by processing TP first. We first demonstrate that the facilitation effect of TP processing on object perception occurs mainly in the M visual system, and then support the subcortical M hypothesis of TP processing by the evidence that the early processing of TP was not affected when cortical function was temporarily damaged by the transcranial magnetic stimulation and when stimuli were biased to M system.

## Introduction

The visual systems of animals are good at rapid detection of threat information in the environment, which is a conserved and innate ability across species. It is commonly considered that visual system has two major separate subdivisions, a magnocellular (M) pathway responsible for fast processing of coarse information, and a parvocellular (P) pathway for slow recognition of details (Callaway, 2005; de Haan and Cowey, 2011; Livingstone and Hubel, 1987; Merigan and Maunsell, 1993). Neuroimaging evidence shows that fast M projections conveying low spatial frequencies facilitate visual object recognition by initiating top-down processes from orbitofrontal to the visual cortex (Bar et al., 2006; Kveraga et al., 2007). However, it is unclear which geometric properties of objects M pathway conveys to accelerate visual object processing. So far, the most commonly exploited functional distinctions between the M and P pathways have been the different sensitivity in spatial and temporal domains, as well as different responses to luminance contrast and color. The M pathway prefers lower spatial frequency and higher temporal frequency, and is highly sensitive to low luminance contrast, while entirely ‘blind’ to chromatic stimuli. The P pathway, by contrast, is more sensitive to stimuli with a higher spatial-frequency but lower temporal-frequency, color-sensitive, but weaker response to low contrast (< 8%) stimuli (de Valois et al., 1977; Field and Chichilnisky, 2007; Goodale and Milner, 1992; Kaplan and Shapley, 1986; Kru, 1979; Lee et al., 1990; Tootell et al., 1988). Meanwhile, the M and P pathways are known as the classical “where” and “what” pathways, which are considered to be mainly located in the dorsal and ventral pathways of the cortex, respectively. Physiological evidence yet demonstrates that the superficial layer of the superior colliculus (SC), a subcortical brain area, shows little color-related activity, and predominantly responses to achromatic information (Schiller et al., 1979; White et al., 2009), which is similar to M function, suggesting that there is probably a subcortical M visual system. As Kveraga et al. (2007) pointed out, a fast subcortical M projection from the thalamus to the prefrontal cortex may be the way the M system facilitates object recognition. In the present study, we proposed a bold hypothesis that M pathway contributes to the fast object recognition by first extracting TP from scenes to build an early object representation. Reason are as follows.

The TP of a figure is the holistic property which remains constant across various smooth shape-changing transformations of an image. For instance, a piece of rubber sheet could change its shape randomly, like, bending twisting, as long as it does not tear, its TP remains the same. But if you drill a hole in the rubber sheet, its TP changes. Thus the number of holes (hereafter referred to as hole) is one of the TP. A topological theory, which holds that the extraction of TP serves as the starting point of object perception, has been proposed to address the fundamental question of what are the primitives of visual perception (Chen, 2005, 1982). This theory has been supported by sufficient evidence across species from insects, rodents to humans, demonstrating that visual system is more sensitive to detect TP differences in images than other non-TP shape differences, and TP is processed automatically and with higher priority (Chen, 1982; Chen et al., 2003; Chien et al., 2012; Han et al., 1999; Huang et al., 2019, 2018, 2011; Todd et al., 1998). Previous evidence from human brain imaging, i.e., functional magnetic resonance imaging (fMRI), however, shows that the inferior temporal cortex (IT) was involved in TP processing (Wang et al., 2007; Zhou et al., 2010; Zhuo et al., 2003). IT is considered to be related to the late stages of processing in the ventral visual stream (Mishkin et al., 1983), thus these neuroimaging findings seem to conflict with the behavioral evidence of preferential processing of TP. Whereupon we come up with a hypothesis that the TP may be processed through a fast subcortical M visual pathway before being projected to the cortical area IT. Our recent fMRI findings provide some support for this subcortical hypothesis, showing that the perception of ‘hole’ was processed in the subcortical regions, i.e., the SC and pulvinar (Meng et al., 2018). Moreover, our latest research on mice provides neurological evidence for TP processing in the SC (Huang et al., 2019).

In the present study, we aimed to investigate whether the TP processing could accelerate object recognition (including TP and other non-TP discrimination) through a fast M pathway, and further identify whether the fast processing of TP occurs in a subcortical M visual system. Stimuli were designed to preferentially engage the M or P systems, i.e., M-biased or P-biased. We adopted an unconscious priming paradigm in Experiment 1 to examine whether the TP processing facilitates object recognition through the M pathway. Then we investigated the sensitivities of the M and P visual systems in processing TP and non-TP in Experiment 2. Further, to test whether a subcortical M visual system is involved in TP processing, in Experiment 3, we temporarily disrupted the function of the primary visual cortex (V1) by applying transcranial magnetic stimulation (TMS) over the occipital lobe at various time delays, and examined the performance for both M- and P-biased stimuli.

## Results

### TP congruence accelerates the recognition of both TP and non-TP through the M pathway

A previous study of unconscious priming found reliable facilitation effect of TP congruence between unconscious prime stimuli and target stimuli on objects’ recognition in both TP and non-TP (orientation) judgements (Huang et al., 2011). Here we combined the unconscious priming paradigm with M/P-biased stimuli (see Figure 1A) to examine whether this TP facilitation effect is specific to the M pathway. The stimuli were either achromatic and of low luminance contrast (< 8% Michelson contrast) to preferentially engage the M system (M stimuli), or isoluminant red-green figures to the P system (P stimuli). The prime/probe stimuli could have one of two possible TP (one hole or no hole), and one of two possible orientations (up or down), yielding four types of prime-probe combinations of TP and orientation, as illustrated in Figure 1B. Participants were requested to report TP (one hole/no hole) or orientation (up/down) of the probe in separate blocks by pressing a response key as soon as possible. There were four separate blocks for combinations of M/P stimuli and two tasks. Different prime-probe congruence pairings were arranged randomly in each block. Participants performed well in all four blocks, with accuracy of 97% ± 0.35% (showing no difference between blocks, F(3,60) = .4, p = .754). The average response times (RTs) of all blocks were submitted into a four-way ANOVA with task, M/P stimuli, TP congruence and orientation congruence as within-subject factors. Significant main effects of task and TP/orientation congruence revealed that mean RTs were shorter for responses to TP than to orientation (F(1,15) = 25.31, p < .001), and RTs were shorter for prime-probe congruent pairings than for incongruent ones, i.e., priming effect (both TP and orientation congruence, Fs(1,15) > 10.694, p < .005). The mean RT for the M stimuli was not different from that for P stimuli (F(1,15) = .199, p = .662). There were significant interactions between task and either type of congruence, between task and M/P stimuli, as well as between M/P stimuli and TP congruence (Fs(1,15) > 6.16, p < .025). These interactions indicated that priming effects were modulated by task and M/P stimuli.

**Figure 1.**
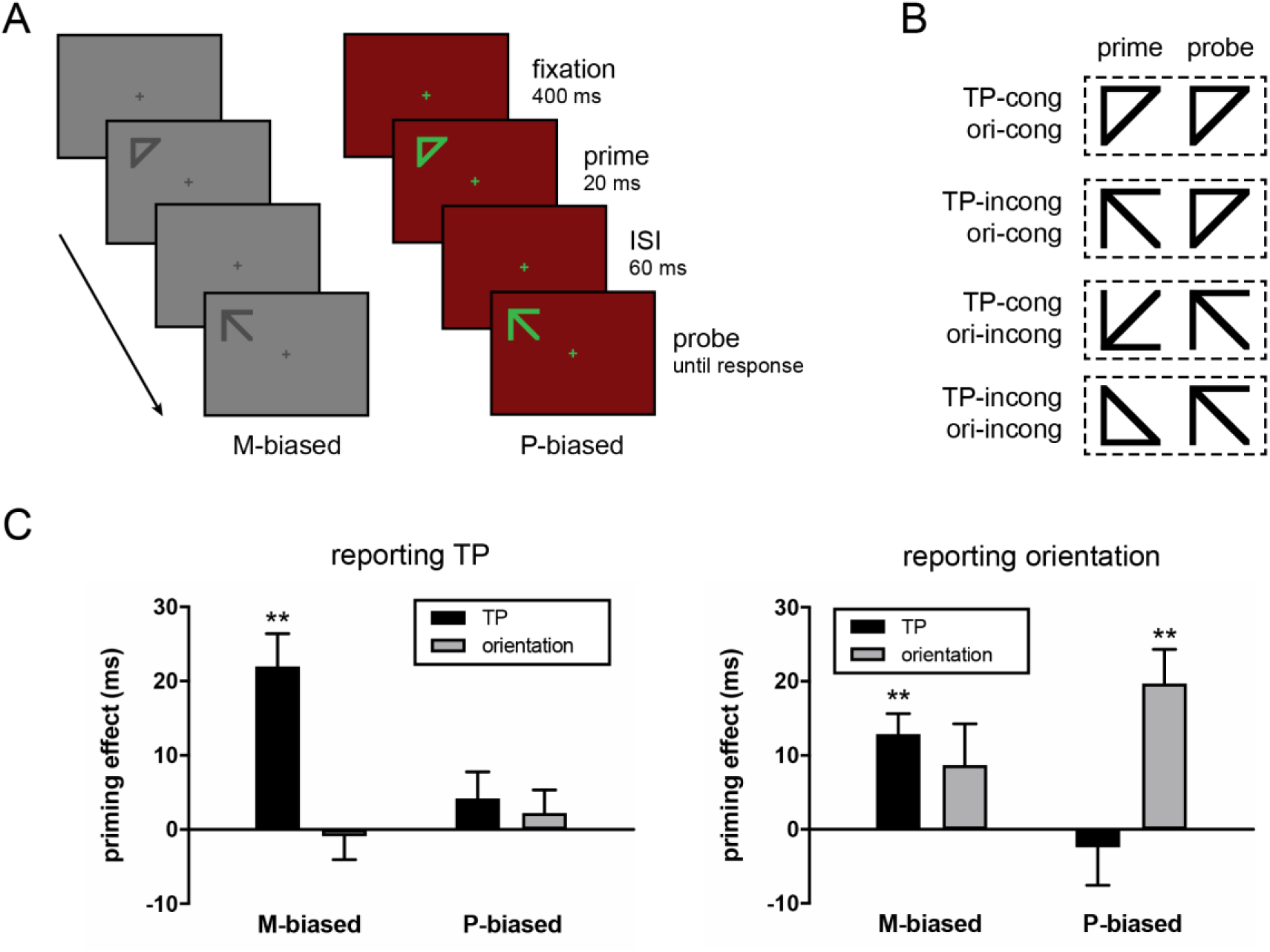
Stimulus procedure, stimulus samples and results in Experiment 1. (A) Stimulus sequence of the priming paradigm under M- and P-biased conditions in Experiment 1. M-biased stimuli were achromatic and of low luminance contrast, while P-biased stimuli were chromatically defined and isoluminant (red-green). The sequence illustrates a prime-probe pair that is incongruent in TP but congruent in orientation. (B) Examples of four prime-probe congruence conditions. The prime and probe could be triangle or arrow, representing two different global topological properties (one hole and no hole), and could have one of the two possible orientations (up or down). Cong = congruent; incong = incongruent; ori = orientation. (C) Priming effects (RT of incongruent minus RT of congruent) of TP and orientation. TP congruence could speed up reports of both TP and orientation, and these priming effects of TP were found only when the stimuli were biased toward M processing, yet not for the P-biased stimuli. The asterisks indicate significant congruence priming effects (** p < .01). Error bars represent the stand errors of the means.

Then two-way ANOVAs with factors of TP congruence and orientation congruence were performed for each of the four conditions (task x M/P stimuli), respectively. There was no significant interaction between two types of congruence for any of the four conditions (Fs(1,15) < 2.407, p >. 142). The priming effects of TP and orientation were shown in Figure 1C, which were calculated by subtracting the RTs of congruence from the RTs of incongruence. We found significant priming effects of TP in both tasks only when the stimuli were biased toward M pathway processing (ts(15) > 4.632, p < .001), yet not for the P-biased stimuli (ts(15) < 1.172, p > .259). In contrast, the priming effect of orientation only occurred when orientation was reported and when stimuli were P-biased (t(15) = 4.249, p < .001). This finding of beneficial priming effects of TP congruence on non-TP (e.g., orientation) recognition consists with Huang’s work (Huang et al., 2011). Critically, we found that this facilitation effect from TP congruence was restricted to the M visual system.

To make sure unconsciousness of the priming effect, we tested the visibility of prime stimuli in another four experimental blocks (2 tasks: TP or orientation x 2 M/P stimuli). The accuracy rates for reporting primes in the four blocks were not different from the chance level of 50% (ts(15) < 1.453, p > .167), indicating that both TP and orientation of the prime stimuli were invisible in M- /P-biased conditions.

### M pathway is more sensitive to TP differences than non-TP shape differences

Experiment 1 adopted RT as an index demonstrating that the TP facilitation effect on object recognition occurs mainly in the M pathway. Our previous study shows that the human visual system is highly sensitive to discriminate topological differences compared to other non-TP differences, which is referred to as TP priority effect (Chen, 1982; Huang et al., 2018). In Experiment 2, we used discrimination sensitivity as another index to investigate whether the TP priority effect shows primarily in the M visual system. As shown in Figure 2A, a stimulus pair was presented for 20 ms, followed by a 100-ms white mask pair. Participants had to judge whether the stimulus pair contains two identical pictures. Geometric figures and letter figures were used as stimuli in Experiments 2A and 2B, respectively. In both experiments, stimuli were either M-biased (luminance contrast < 8% Michelson contrast) or P-biased (isoluminant red-green). The luminance contrast of M-biased achromatic stimuli, and the color contrast of P-biased red/green stimuli were slightly adjusted to keep an overall mean average accuracy of about 70%. Stimuli were divided into two types according to their TP: one-hole and no-hole figures (see Figure 2B). Therefore there were two types of different stimulus pair, i.e., TP different and non-TP different. The TP different and non-TP different trials were arranged in separate blocks, in each of which the same and different pairs were evenly and randomly distributed. The average accuracy in discriminating letter figures and geometric figures was about 71%. D prime was calculated as an index of discrimination performance and submitted into a two-way repeated ANOVA with M/P stimuli and discrimination type (TP or non-TP different) as factors. For both geometric and letter stimuli, we found a significant main effect of discrimination type (Fs > 5.013, p < .04) and significant interaction (Fs > 5.42, p < .029), but no significant effects of M/P stimuli (Fs < 1.267, p > .271). Further post-hoc analyses revealed that only when the stimuli were M-biased was there a significant advantage of TP-difference discrimination over non-TP-difference discrimination, i.e., TP priority effect (for both letter and geometric stimuli, ts > 3.894, p < 0.001). This effect disappeared in the case with P-biased stimuli (both ts < 1.385, p > .179, see Figure 2C). These findings indicated that the M visual pathway was more sensitive in discriminating TP than non-TP shape differences, while the P pathway exhibited no preference.

**Figure 2.**
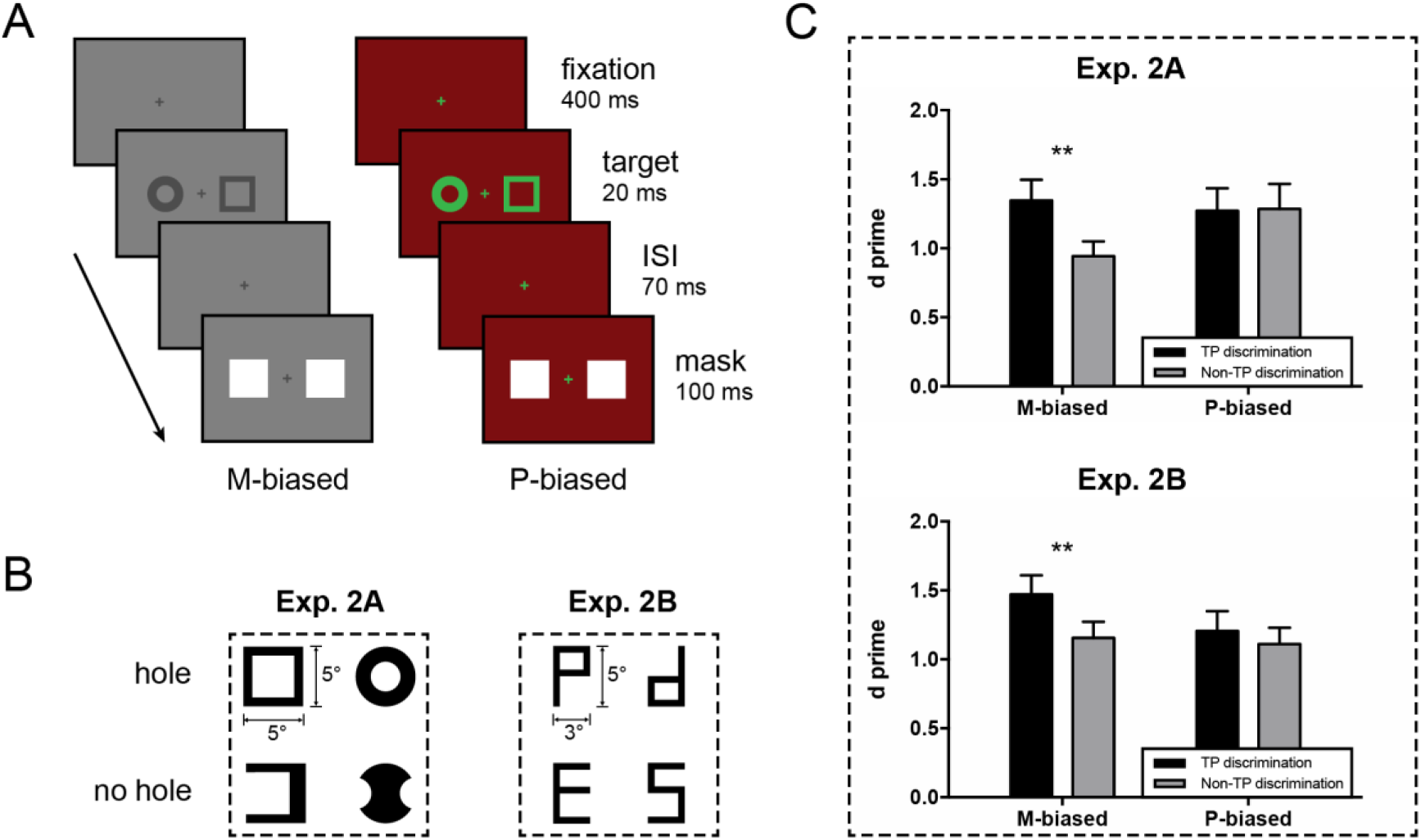
Stimulus procedure, stimulus samples and results in Experiment 2. (A) Discrimination paradigm under M- and P-biased conditions in Experiment 2. The stimulus pair shown was a sample of non-TP-different discrimination. (B) Geometric and letter figures were used as stimuli in Experiment 2A and 2B, respectively. The figures were divided into two types according to their TP (hole vs. no hole). (C) Performance (d prime) in discriminating TP and non-TP differences with M- and P-biased stimuli in Experiment 2A and 2B. Results indicated that the M visual pathway was more sensitive in discriminating TP than non-TP shape differences, while the P pathway exhibited no preference. The asterisks indicate a significant difference between TP and non-TP discriminations (** p < .01). Error bars represent the stand errors of the means.

### Subcortical M visual system is engaged in the early processing of TP

Previous TMS work has demonstrated that TP might be processed through a rapid subcortical visual pathway bypassing V1 (Du et al., 2011). Moreover, results from Experiments 1 and 2 showed that the fast processing of TP primarily occurs in the M pathway. Thereupon, in Experiment 3 we tested whether a subcortical M system was responsible for the fast TP processing. If the early processing of TP occurs in a subcortical system, it is not expected to be affected when the cortical function is temporarily damaged by the TMS. Stimuli were biased to the processing of M or P pathway (see Figure 3A), and consisted of four items, either all the same or one different from the other three. Participants were instructed to report whether the four items were the same. As shown in Figure 3C, there were three types of stimuli with items differing either in non-TP (Experiment 3A,) or in TP (Experiment 3B and 3C). It is worth noting that we used an s-like stimulus which was used in previous studies and was specially designed to match the ring in both stimulus area and spatial frequencies (Chen et al., 2003; Huang et al., 2018). TMS pulses were applied to disrupt V1 function at varies of temporal delays after stimuli onset. The TMS intensity was set at 100% of the phosphene threshold for each participant.

**Figure 3.**
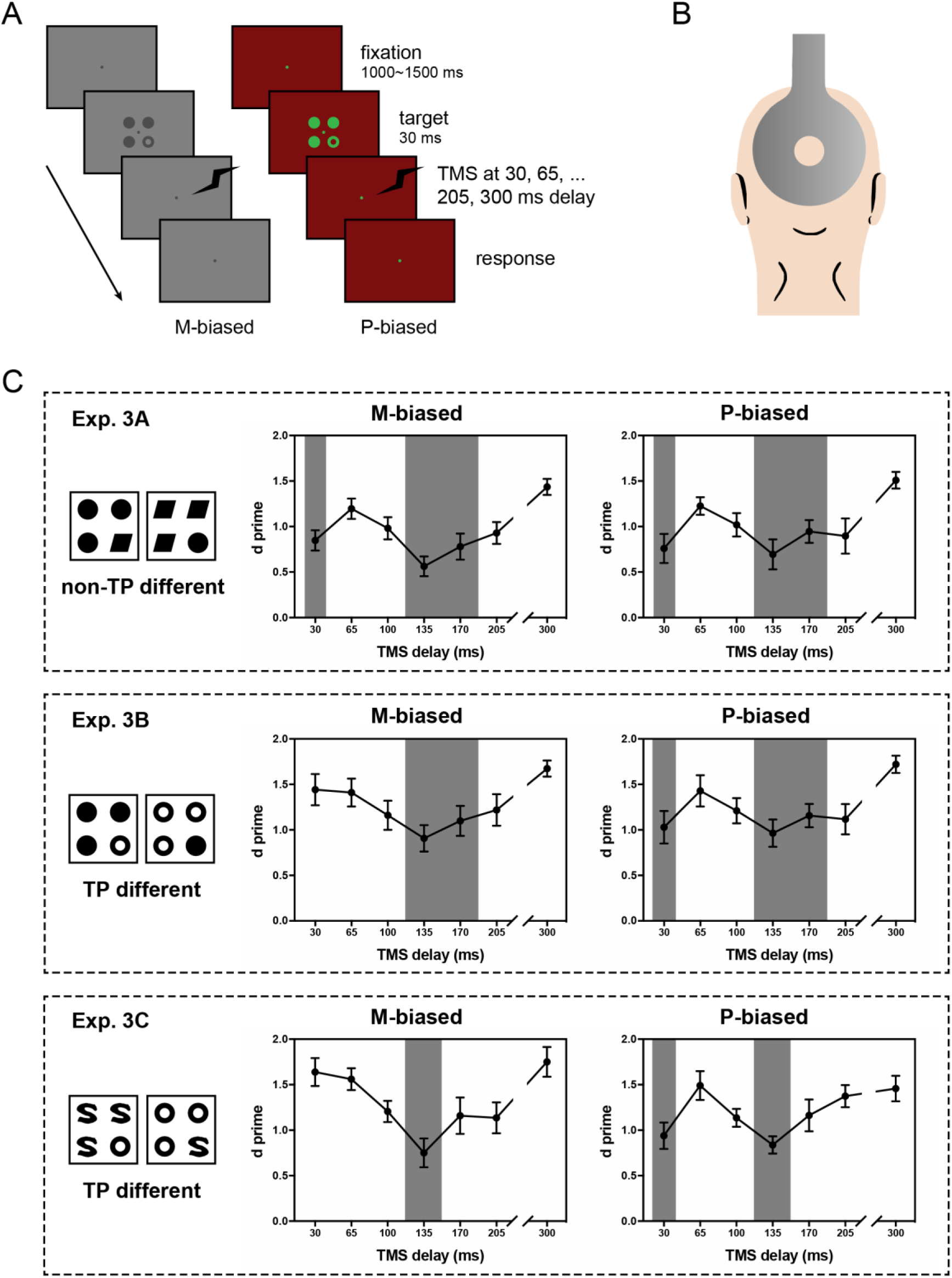
Stimulus procedure, TMS coil and results of Experiment 3. (A) Stimulus sequence of discrimination paradigm under M- and P-biased conditions in Experiment 3. The M- /P-biased stimuli designs were identical to those in Experiments 1 and 2. The target here illustrates TP-difference discrimination. (B) Illustration of the TMS coil position. Here the coil was positioned over the occipital pole, with the lower rim of the coil 1 ∼ 2 cm above the inion, to target primary visual cortex (V1). (C) Performance (d prime) in discriminating TP and non-TP difference in Experiment 3. The grey columns indicate significant decline of d prime (p < .05) as compared with the baseline level, which was defined as the average d prime at 205 ms and 300 ms delays.

D prime values for Experiments 3A, B and C were entered into two-way ANOVAs with SOA (TMS delays) and M/P stimuli as factors, respectively. For each experiment, the main effect of SOA was significant (Fs > 3.209, p < .019) while the main effect of M/P stimuli was not significant (Fs < .874, p > .365), indicating that TMS influenced performance and similar task difficulty of the M and P stimuli were matched. Importantly, the interaction was significant for TP-different stimuli (Fs > 2.995, p < .025), suggesting that SOA has different effects with M or P stimuli. In contrast, there was no significant interaction for non-TP-different stimuli (Exp. 3A, F(6,90) = .393, p = .813). Specifically, when discriminating non-TP different stimuli (both M- and P-biased), the d prime values were significantly below the baseline level (defined as the average performance at the 210 ms and 300 ms delays) at the TMS delays of 30 ms, 135 ms and 170 ms (ts > 2.233, p < .041). The first temporal phase (30 ms) happens in the early time of visual processing, and usually reflects the feedforward processing of visual information travelling from the retina to V1, while the latter temporal phase (135 ∼ 170 ms) reflects the feedback modulation of higher cortical areas to V1 (Bullier, 2001; Gilbert and Li, 2013; Hupé et al., 1998; Lamme and Roelfsema, 2000). The TMS disruption of V1 significantly reduced the performance of discriminating non-TP in both feedforward and feedback temporal phases of visual processing. The d prime values for TP-different stimuli in Experiment 3B and 3C were separated into M and P stimulus groups. Simple effect analyses showed a significant effect of SOA for both M and P stimuli in Experiment 3B and 3C (Fs > 2.724, p < .017). Further post-hoc analyses revealed that for the P-biased stimuli there was a significant drop in performance at an early phase (30 ms, ts > 3.8, p < .002) and a late phase (Exp. 3B, 135 ∼ 170 ms, ts > 2.342, p < .033; Exp. 3C, 135 ms, t(15) = 4.927, p < .001). For the M-biased stimuli, there was also a drop at the late temporal phase remained (Exp. 3B, 135 ∼ 170 ms, ts > 2.61, p < .02; Exp. 3C, 135 ms, t(15) = 3.284, p = .005), whereas the early drop at 30 ms disappeared (Exp. 3B and 3C, Fs < 1.279, p > .22), suggesting that the disruption of V1 only affected the performance of discriminating TP with M-biased stimuli at the late temporal phase, not at the early phase. These results were in agreement with the previous finding that the early processing of TP was not affected by the TMS (Du et al., 2011). More importantly, the performance of discriminating TP-different stimuli was found significantly declined at the early temporal phase only when the stimuli were P-biased rather than M-biased, suggesting that M-biased stimuli could be early processed through a subcortical pathway.

## Discussion

The present study aimed to test our hypothesis that the subcortical M visual system may be engaged in the fast processing of TP. Specially-designed stimuli were used to bias visual processing toward either M or P pathway. There are three main findings, including (1) the facilitation effect of TP processing on object perception occurs mainly in the M visual system; (2) the visual system is more sensitive to detect TP differences than non-TP differences between images, and this TP priority effect in perception relied on the processing of M pathway; (3) the TP of objects may get early processing in a subcortical magnocellular system, not through the cortical visual pathway.

The first finding is consistent with our previous work (Huang et al., 2011) on the facilitation effect of unconscious topological processing on objects’ feature recognition. Moreover, this TP facilitation effect critically depended on the processing of M pathway, and disappeared when the M pathway was inhibited (i.e., P-biased condition). The finding indicated that the visual system facilitates object recognition by preprocessing TP in the fast M pathway, as evidenced by the shorter RTs of TP. According to the topological theory proposed by Chen (Chen, 2005, 1982), extraction of TP is used to construct object representation. When a prime stimulus appears, an initial object representation based on TP has been established. If the following probe stimulus is of the same TP, the existing object representation will be used, however, if not, there will be a new object representation to be constructed, which will cost extra reaction times regardless of what property is to report.

In addition to the evidence from response speed, our second finding from discrimination sensitivity experiment provided further evidence for the specificity of M visual system in processing TP difference between figures. It is worth noting that the possible confounding factors, like spatial frequencies and stimulus area (luminance), were excluded in our experiments, as these factors are distinguishing features between M and P visual systems (Stuart et al., 2012). Stimuli in each of the three experiments were matched in stimulus area, except the stimuli in Experiment 3b. We further conducted spatial frequency analyses of stimuli figures. Figure 4 showed the power spectra (2D Fourier transformation) results and the differences in power spectra (dps, defined as Sum [(X-Y)^2^], see Chen et al., 2003) between figures. The dps values of TP different stimulus pairs (e.g., ‘S’ and ‘O’, dps = 3.52) are much smaller than those of TP same stimulus pairs (e.g., disk and parallelogram, dps = 25.18). Thus, it is TP rather than spatial frequencies or luminance that M pathway extracts from scenes for fast object recognition.

**Figure 4.**
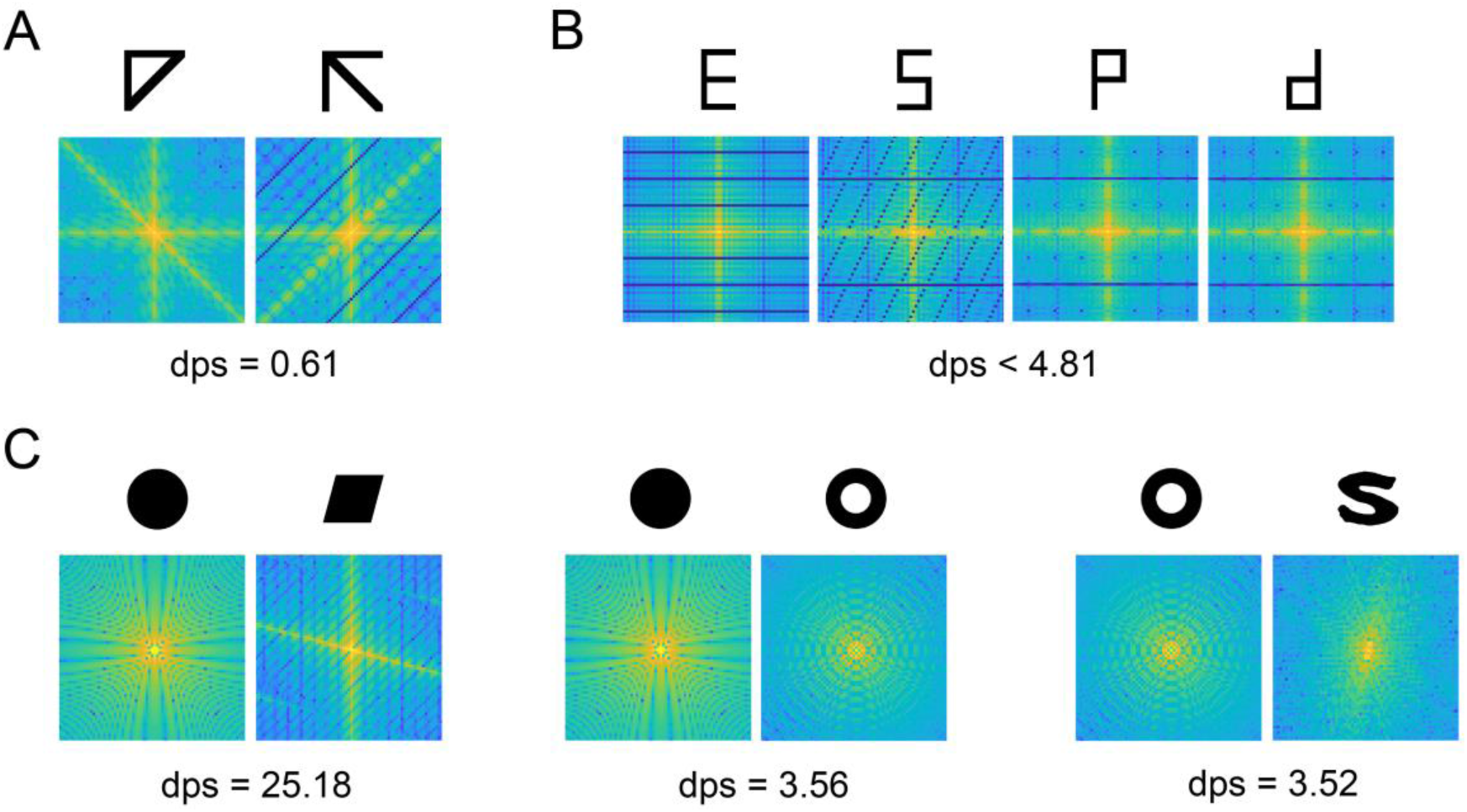
Spatial frequency analyses of stimuli figures. (A) The power spectra (2D Fourier transformation) of the arrow and triangle used as stimuli in Experiment 1. The differences in power spectra (dps, defined as Sum [(X-Y)^2^]) between the two figures are 0.61. (B) The power spectra of the letter figures used in Experiment 2b. The dps between the four figures are smaller than 4.81. (C) The power spectra of stimuli in Experiment 3. The dps values between TP-different stimuli (about 3.5) were much smaller than that between TP-same stimuli (25.18). These results suggest that it is TP rather than spatial frequency that the M pathway extracts for fast object recognition.

The findings from Experiments 1 and 2 indicate that the M pathway could be responsible for the TP processing. The dorsal visual stream, known as the classical M pathway, is responsible for direction and motion perception, whereas there is no evidence from neuroimaging showing that TP is processed in dorsal visual cortex (Wang et al., 2007; Zhou et al., 2010; Zhuo et al., 2003). On the contrary, evidence from human fMRI and mouse neurological studies supports for the involvement of some subcortical regions, e.g., SC and pulvinar, in the TP perception (Meng et al., 2018). The present TMS results provide further evidence for the subcortical processing of TP and the existence of the subcortical magnocellular visual system. Likewise, Zhang and his colleagues have found that the superficial layer of the SC has similar spatiotemporal and contrast responses as the magnocellular layers of the lateral geniculate nucleus, which then projects to the V1 (Zhang et al., 2015). Kveraga et al. have also suggested that such a subcortical M pathway may be responsible for the fast object recognition, based on their findings of a greater activation for M-biased stimuli than P-biased stimuli in the amygdala, which is also a pivotal region in the subcortical system (Kveraga et al., 2007). The subcortical visual system is considered to stem from the SC, which receives direct input from the retinal ganglion cells (Perry and Cowey, 1984). A subcortical visual pathway, which may include the SC, pulvinar and amygdala, is considered to be responsible for the innate defensive behavior (Morris et al., 1999; Wei et al., 2015). For example, when mice are presented with a threatening looming stimulus, they demonstrate defensive flight-to-nest behavior. Interestingly, our recent work shows that the topological change in a looming stimulus weakened the defensive responses and reduced cell activation in the SC, indicating that detection of threatening information is dependent on TP-based object representations and the processing of TP and threatening information may commonly occur in the SC (Huang et al., 2019). The present findings provide human evidence from TMS study for the involvement of a subcortical M pathway in TP processing, which could be a newly discovered function of the conservative subcortical visual system.

Exploring the function of M pathway is not only an important research topic in vision, but also sheds light on the pathogenesis of magnocellular-dysfunction related brain disorders, such as schizophrenia, autism, dyslexia and glaucoma (Butler et al., 2005; Eden et al., 1996; Greenaway et al., 2013; Shabana et al., 2003; Zhang et al., 2015). Further work identifying and characterizing neural circuits involved in subcortical topological perception may help in the understanding and treatments of these disorders.

## Materials and Methods

### Participants

One hundred and six human participants were recruited in the study, with 16 in Experiment 1 (6 males, average age 22 years), 17 in Experiment 2A (8 males, average age 23 years), 25 in Experiment 2B (9 males, average age 22 years), 16 in Experiment 3A (9 males, average age 22 years), 16 in Experiment 3B (7 males, average age 23 years) and 16 in Experiment 3C (7 males, average age 20 years). All participants were right-handed, and reported normal or corrected-to-normal vision, including intact red-green color vision. Participants provided written informed consent, and were paid to compensate their time. The study was performed in accordance with the Declaration of Helsinki, and was approved by the ethics committee of Institute of Biophysics, Chinese Academy of Sciences.

### Apparatus

Visual stimuli were presented on a 19-inch CRT monitor with 100-Hz refresh rate and 1024 × 768 pixel resolution. The luminance of the monitor was calibrated (gamma correction) with a photometer. A Magstim 200 (Whiland, U.K.) stimulator with a 140-mm circular coil was used in the TMS study. Stimulus presentation and data acquisition were controlled by custom programs written in Matlab (Mathworks, Natick, USA) using Psychtoolbox (Brainard, 1997).

### Stimuli and Procedures

Participants were required to maintain their fixation throughout a trial, with their heads stabilized in a chin rest at the viewing distance of 57 cm in a dark room. In all experiments, both M-biased and P-biased stimuli were adopted to bias visual processing toward M and P pathways respectively, and then tested in separate blocks with the same procedures. The M-biased stimuli were achromatic gray-scale figures with low luminance contrast (< 8% Michelson contrast), and the P-biased stimuli were chromatically defined, with isoluminant red/green as the colors of background and stimuli, respectively. The luminance of the isoluminant red/green colors were obtained through a minimal flicker procedure to match subjective brightness, which was used in previous studies (e.g., Zhang et al., 2016).

Line drawings of arrows and triangles that matched in the area and spatial frequencies were used as stimuli in Experiment 1 (see Figure 1A). The prime (1.13°) and probe (1.88°) stimuli were presented at one of four quadrants of the screen with their centers at an eccentricity of 5.6°. Each trial consisted of a 20-ms prime stimulus, a 60-ms blank screen, and a probe stimulus presented until the participant’s response. Participants were requested to report TP (one hole/no hole) or orientation (up/down) of the probe in separate blocks by pressing a response key as soon as possible. The RT in each trial was calculated from the onset of the probe stimulus and recorded using a parallel port response keypad. The inter-trial interval ranged randomly from 0.5 s to 1.5 s. Each participant completed 768 trials in total, with 192 trials in each of the four blocks (2 tasks: TP or orientation x 2 M/P stimuli: M- or P-biased) and 48 trials for each condition (2 TP congruence x 2 orientation congruence).

In Experiment 2, stimulus pairs were presented at 6° to the left and right of screen center, followed by a pair of white masks (each 8° × 8°), as illustrated in Figure 2A. The stimulus pairs were either chosen from four letter figures (‘E’, ‘S’, ‘P’, ‘d’), or four geometric figures (see Figure 2B). These figures can be divided into two types according to their TP (hole). For instance, as shown in Figure 2B, ‘P’ (or ‘d’, with one hole) are topologically different from ‘E’ (or ‘S’, with no hole), on the other hand, ‘P’ and ‘d’ (or ‘E’ and ‘S’) are of the same TP but different in shape (referred to as ‘non-TP different’). Both the letter figures (3° × 5°) and geometric figures (5° × 5°) matched in the stimulus area (luminance), and the letter figures also matched in spatial frequencies. Participants had to make a two-alternative forced choice (2AFC) and judge whether a stimulus pair consisted of the same figure (Figure 2A). Each trial began with a stimulus pair presented for 20 ms, followed by a 100-ms white mask pair, with a 70-ms ISI in between. Participants gave their answers by pressing a response key without time limit. The TP different and non-TP different trials were arranged in separate blocks with block order balanced across participants. In each block, the same and different trials were of the same proportion (50%). The luminance contrast of M-biased achromatic stimuli, and the color contrast of P-biased red/green stimuli were slightly adjusted for each participant to keep an overall mean average accuracy of about 70%. Each participant finished 384 trials, with 96 trials in each of the four blocks (2 discrimination types: TP or non-TP different x 2 M/P stimuli: M- or P-biased).

In Experiment 3, three types of stimuli were tested separately (non-TP difference discrimination in Experiment 3A and TP difference discrimination in Experiment 3B and 3C). Each stimulus consisted of four items and were presented in a 2° × 2° region in the screen center. The four items could be all the same, or with one item different from the rest either in projective properties (non-TP different, e.g., disk vs. parallelogram) or in ‘hole’ (TP different, e.g., disk vs. ring, and ring vs. s-like figure), as illustrated in Figure 3C. The disk and ring extended 0.64° × 0.64° of visual angle, and the parallelogram matched the disk in the stimulus area. The s-like figure was specially designed to match the ring in both stimulus area and spatial frequencies. Similar to Experiment 2, M-biased and P-biased stimuli were tested in separate blocks and stimuli were adjusted within a small range for each participant to control average accuracy of about 70%. TMS pulses were applied over the primary visual cortex in the discrimination task. Phosphene thresholds, i.e., minimal TMS intensity for inducing a visual sensation, were measured for each participant after dark adaptation. As illustrated in Figure 3B, the lower rim of the circular coil was initially positioned 1.5 cm above the inion (over the occipital pole), and then adjusted until the visual phosphene was induced in the central visual field. The TMS intensity was set at 100% of the phosphene threshold for each participant. The average TMS intensities were about 61% of maximum output. During each trial, a stimulus array was presented for 30 ms, followed by a single-pulse TMS pulse delivered at various delays after stimulus onset (TMS delay: 30, 65, 100, 135, 170, 205 and 300 ms). Participants were instructed to report whether the four items were the same without time limit. Each participant completed 448 trials in total, with 224 trials for each M- / P-biased sessions, and 32 trials for each TMS delay. The baseline level was defined as the mean performance on the trials in which TMS pulses were delivered between 200 and 300 ms after stimuli onset (Jolij and Lamme, 2005).

## Acknowledgement

This work was supported in part by the Ministry of Science and Technology of China grant (2015CB351701), the National Nature Science Foundation of China grant (31730039), the Chinese Academy of Sciences grant (XDBS32000000), the Guangdong Key Lab of Brain Connectome (2017B030301017), and Shenzhen Science and Technology Research Funding Program (JCYJ20170818161400180).

